# Competitive exclusion, experience-based learning, and human fishing, influence activity patterns in Juvenile and Adult Brown Pelicans (*Pelecanus occidentalis*)

**DOI:** 10.1101/2024.10.01.616051

**Authors:** Dennys Plazas-Cardona, Juan Camilo Ríos-Orjuela, Juan D. Wilches-Vega

**Author notes:** Una versión en español de este manuscrito se encuentra disponible en: dx.doi.org/10.6084/m9.figshare.27144603.

## Abstract

The activity patterns of young and adult individuals offer insights into exclusion and adaptation dynamics in seabirds. Juveniles often show differences in their daily behaviors compared to adults, especially in activities like feeding. We focus in two contrasting hypotheses to explain the age-related differences in activity patterns of Brown Pelicans (*Pelecanus occidentalis*): the “competitive exclusion” hypothesis and the “experience-based adaptation” hypothesis. The competitive exclusion hypothesis posits that adult pelicans, with superior foraging skills, actively exclude juveniles from prime feeding areas, leading to temporal segregation. In contrast, the experience-based adaptation hypothesis suggests that juveniles exhibit different activity patterns as they undergo a learning process, gradually improving their foraging efficiency through trial and error. Using continuous focal sampling, we analyzed the activity peaks of both juveniles and adults, revealing significant temporal segregation between age groups. Additionally, the impact of human fishing activity on pelican behavior was assessed, showing alterations in natural foraging patterns. These findings contribute to the understanding of resource use and competition dynamics in seabirds and highlight the importance of considering both biological and anthropogenic factors when analyzing seabird behavior.

## Introduction

Activity patterns in individuals are a feature of behavioral biology that can be influenced by various factors, including biological and environmental aspects (Waugh et al. 2018, Serratosa et al. 2020). Understanding how these patterns differ between age groups provides critical insights into their life history strategies and the mechanisms of resource use and competition. Age-related differences in activity are common in many species, where juveniles often exhibit distinct behavioral patterns compared to adults. These differences can have far-reaching consequences on their foraging efficiency, survival rates, and overall fitness (Orians 1969, Gochfeld and Burger 1984, Marchetti and Price 1989).

In seabird populations, competition for food resources is a major factor influencing activity patterns. Juveniles, who are often less experienced and less efficient foragers, may be excluded from prime foraging sites by more dominant adults (Riotte-Lambert and Weimerskirch 2013). These exclusionary behaviors are not uncommon and have been documented in a variety of seabird species, where adults secure access to key resources, leaving juveniles to forage in suboptimal conditions (Riotte-Lambert and Weimerskirch 2013, Bodey et al. 2014, Afán et al. 2019). In the case of Brown Pelicans, foraging primarily involves plunge diving to capture fish, a skill that improves with age. Adults tend to exhibit higher foraging efficiency and success rates than juveniles, which contributes to disparities in resource acquisition between the two groups (Orians 1969).

One widely studied condition in behavioral research is the idea that young individuals behave differently from adults throughout the day, and some of the hypothesis around the interactions between adults and juveniles in sea birds posits that adults actively exclude juveniles from engaging in activities such as feeding, perching, comfort behaviors, and flight, thereby securing more resources for themselves (Yoda et al. 2011, Corbeau et al. 2020). However, some other evidence suggest that juveniles could inherently avoid sharing these activities with adults due to their inability to compete effectively for resources (Riotte-Lambert and Weimerskirch 2013, Pettex et al. 2019).

Here, we focus on two contrasting (but not exclusive) hypotheses; the first one is the “competitive exclusion” hypothesis, which, in our case, posits that adult pelicans, due to their superior foraging skills and territorial dominance, actively exclude juveniles from prime feeding areas and perch sites. This behavior would result in temporal and spatial segregation, with adults monopolizing resources during peak feeding times, forcing juveniles to adjust their activity to non-peak times or less productive areas (Rescigno and Richardson 1965).

The second hypothesis is the “experience-based adaptation” hypothesis, which suggests that the observed differences in activity patterns are largely due to the developmental learning process that juveniles undergo. According to this hypothesis, juveniles’ less efficient foraging strategies lead them to adopt different temporal activity patterns as they gradually improve their skills through trial and error. Over time, their foraging behaviors will begin to resemble those of adults, but in the early stages of life, they must compensate for their lack of experience by spending more time searching for food and flying longer distances (Pyke 1984, Dall et al. 2005).

Evidence supporting the first hypothesis can be observed in the foraging behaviors of juvenile seabirds, which often differ significantly from those of adults (Riotte-Lambert and Weimerskirch 2013, Pettex et al. 2019). Studies indicate that juvenile seabirds exhibit inferior foraging skills compared to adults, leading to lower feeding success and increased mortality rates (Riotte-Lambert and Weimerskirch 2013). This discrepancy can drive adults to exclude juveniles from prime foraging areas, as evidenced by the higher foraging site fidelity and efficiency seen in adults (Grecian et al. 2018).

Conversely, the second hypothesis is supported by observations of juvenile seabirds adopting different movement and foraging strategies to avoid competition with adults (Porter and Sealy 1982, Mendez et al. 2017, Pettex et al. 2019). Juveniles are often found to utilize different areas or engage in less competitive behaviors to minimize direct interactions with adults. For example, juvenile wandering albatrosses exhibit distinct movement patterns, covering shorter daily distances and spending more time resting compared to adults, suggesting avoidance of direct competition (de Grissac et al. 2016). Additionally, juvenile seabirds could have higher mortality rates due to their less efficient foraging skills and greater susceptibility to predation and human activities (Waugh et al. 2018, Afán et al. 2019).

Research indicates that juveniles gradually develop more efficient foraging techniques and movement strategies through a process of trial and error and learning from older, more experienced individuals. For example, studies have shown that juvenile wandering albatrosses initially exhibit less efficient foraging patterns and lower flight efficiency compared to adults (Riotte-Lambert and Weimerskirch 2013, Corbeau et al. 2020). However, over time, they progressively improve these skills, aligning their behaviors more closely with those of experienced adults (Weimerskirch et al., 2006). This learning period is crucial for the survival and eventual reproductive success of juveniles, as it allows them to adapt to the environmental conditions and resource availability in their habitats (Grecian et al. 2018).

The Brown Pelican (*Pelecanus occidentalis*) serves as an exemplary model for studying activity patterns and inter-age group interactions in marine birds due to its distinct biological and behavioral traits (Orians 1969, Walter et al. 2013). Found along coastal regions of North and South America, this species exhibits diverse foraging strategies and habitat use that are essential for understanding the dynamics between adults and juveniles. Brown Pelicans are known for their plunge-diving technique to capture fish, a skill that improves with age and experience. Studies have shown that adult pelicans have higher foraging success compared to juveniles, indicating significant differences in skill and efficiency (Orians 1969). This variation provides a clear basis to investigate whether juveniles avoid competition with adults due to inferior foraging abilities or if adults actively exclude juveniles from prime foraging locations.

Human activities, particularly fishing, have a profound impact on the behavior and activity patterns of seabirds, including the Brown Pelican (*Pelecanus occidentalis*). Fishing practices, such as discarding bycatch, can create readily accessible food sources that alter the natural foraging behaviors and movement patterns of these birds. For instance, research has shown that seabirds exploit fishery discards, which can lead to changes in their spatial distribution and foraging strategies, making them more site-faithful to areas with consistent discard availability (Bartumeus et al. 2010, Waugh et al. 2018). In the case of the Brown Pelican, human activities such as fishing and coastal development can disrupt their natural behaviors and habitat use. Pelicans have been observed to alter their foraging locations and times in response to the presence of fishing boats and human recreational activities. These disturbances can lead to increased energy expenditure as birds are forced to relocate to less optimal foraging sites, potentially affecting their overall fitness and reproductive success (Lamb et al. 2018). Such behavioral modifications highlight the importance of managing human activities to mitigate their impact on seabird populations and ensure the conservation of species like the Brown Pelican.

The primary objective of this research is to evaluate the activity patterns of juvenile and adult Brown Pelicans in a natural setting and to determine the extent to which competition and experience drive these patterns. Additionally, this study seeks to assess the influence of human activity on these behaviors, particularly in a region where fishing practices are prevalent and may provide anthropogenic food sources.

## Methods

### Study site

The study was conducted in El Faro, Cabo de la Vela, located in La Guajira, Colombia (12°12′24″N, 72°10′24″O; Fig 1). This area is characterized by a hot and dry climate, with an average annual temperature of 28.2°C. The maximum elevation is 47 meters above sea level and annual precipitation ranges between 354 and 1170 mm. The region is known for its constant winds, high evaporation rates, and well-defined seasons. Also, there is a tropical upwelling system in the area, which significantly enhances the availability of marine resources. The upwelling brings nutrient-rich waters to the surface, promoting high productivity and an abundance of small pelagic fish, a primary food source for Brown Pelicans (Paramo et al. 2003). The area’s sea surface temperature and salinity, modulated by seasonal upwelling, provide a diverse and dynamic habitat that supports a rich marine ecosystem (Criales-Hernández et al. 2006). The coastal environment of La Guajira also includes a variety of benthic habitats, such as sandy bottoms and coral reefs, further enhancing its ecological complexity and resource availability (Chasqui et al. 2013). These conditions make Cabo de la Vela an excellent natural laboratory to investigate the interactions between adult and juvenile pelicans, examining hypotheses related to competition and resource partitioning within a highly productive marine environment.

**Figure 1.**
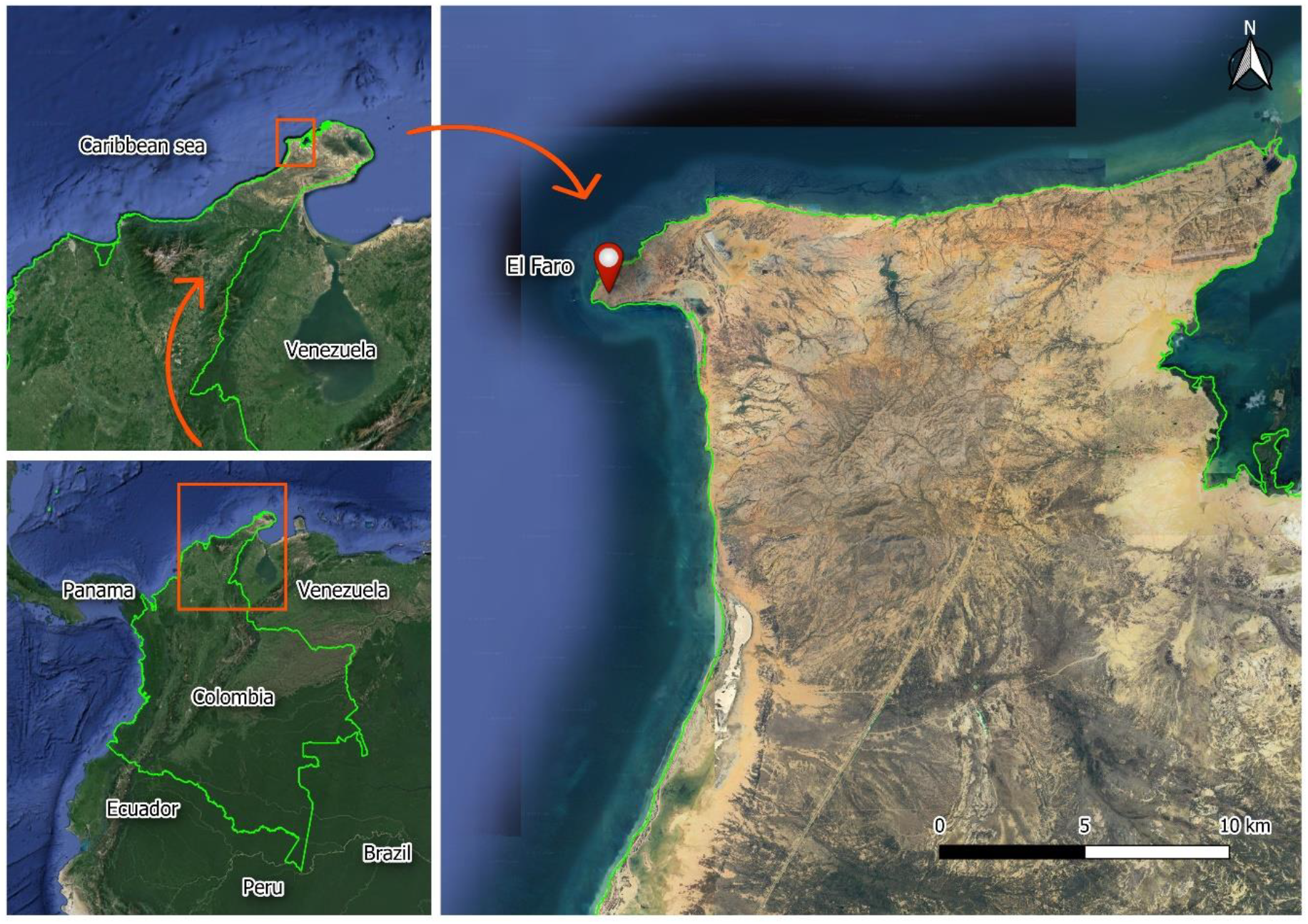
Map of study site. El Faro, Cabo de la Vela, La Guajira, Colombia (12°12′24″N, 72°10′24″O).

### Data collection

This study was conducted from March 11 to 18, 2013. We employed the Continuous Focal Sampling method (Lehner 1992, Rose 2000), observing daily from 5:30 h to 18:30 h, and covering 3 km each day. Our objective was to document the activity peaks of Brown pelicans in relation to individual developmental stages and anthropogenic influences. Juvenile Brown pelicans were identified using plumage color (Walter et al. 2013). Activities were categorized, and each occurrence of an activity was recorded as an observation event.

The activities performed by the birds were classified into four categories: comfort, rest, feeding, and flight, according to Amat (1984). The comfort category includes any activity that provides comfort and maintenance to the bird, such as the “cross position” where the wings are partially extended, and preening, which involves any grooming of feathers or appendages. The rest category refers to periods when the bird is stationary, either individual or group and can occur on water or land. Feeding involves the acquisition and ingestion of food, including fishing and disputes, which involve stealing food from the beak, potentially occurring between individuals of the same or different species. The flight category includes all activities involving wing movement for traveling between two points. This behavior can be individual or group.

We also documented the hours of human fishing activity throughout the day to assess its potential impact on pelican activity peaks.

### Statistical analysis

We used the R package “circular” (Agostinelli and Lund 2022) to transform the data into a circular format based on a 24-hour time scale. After transforming the data, we calculated descriptive statistics, including the mean and standard deviation, to summarize the central tendency and variability of the circular data. Additionally, we computed the circular concentration coefficient, rho (ρ), both for the complete dataset and for each activity type. The rho (ρ) statistic measures the concentration or dispersion of circular data and is used to assess the circular association between observations. Technically, ρ reflects the strength of the circular correlation, with its value linked to the resultant vector’s mean. A ρ value close to 1 indicates that the data points are tightly concentrated around a mean direction, whereas a value near 0 signifies high dispersion and little directional consistency among the data.

To compare whether the circular data for adults and juveniles (i.e., angular measures or cyclic times) originated from the same distribution, we conducted Watson’s Two-Sample Test of Homogeneity (Zar 1976). This test was applied both to the overall dataset and to each individual activity type, allowing us to determine whether the distributions between the two groups were statistically similar or significantly different.

A heatmap was generated to visually assess the correlation between activity type and time of day (start time) for different age groups (adults and juveniles). The heatmap represents the intensity of activity through color gradients, where the value of each cell corresponds to the frequency or magnitude of the activity at a given time. This approach allows for easy identification of patterns, such as periods of higher or lower activity throughout the day. The heatmap was constructed using R’s “ggplot2” and “Reshape2” packages (Wickham 2007, 2016).

We used density plots to visualize the distribution of activity occurrences throughout the day for both adults and juveniles across different activity types. These plots provided a smoothed estimate of the frequency of observations over time. We adjusted the transparency of the density curves to ensure a clear comparison between adults and juveniles.

We employed raincloud plots to visualize the distribution of activities over time by combining violin plots, boxplots, and jittered data points. This method allowed us to simultaneously present the overall data distribution (via violin plots), statistical summaries (via boxplots), and individual data points (via jittered points), providing a comprehensive view of the data. All plots were generated using the “ggplot2” (Wickham 2016), “ggdist” (Kay 2023) and “reshape2” (Wickham 2007) packages in R.

In addition to the circular data analysis, we use raw data to assess the time and activity patterns between adults and juveniles. The analysis was performed to compare two main aspects: the distribution of activity times (it means the time of day used per activity, also mentioned as start times) and the cumulative time spent on each activity.

We applied the Shapiro-Wilk test to assess the data normality of the activity start times for both adults and juveniles. As the data were not normally distributed (p-value < 0.05 for all groups and activities), we proceeded with the Mann-Whitney U test to determine if there were significant differences in the times at which adults and juveniles engaged in various activities. The Mann-Whitney U test was also applied to compare start times for specific activities.

In terms of cumulative activity time, we aggregated the data by counting the number of individuals engaged in each activity throughout the day. We again checked for normality using the Shapiro-Wilk test and applied the Mann-Whitney U test when the data did not meet normality assumptions. However, because cumulative time had only one value per group for each activity (lacking variability), we performed a categorical analysis of Fisher’s exact test (Warner 2013) on a contingency table that summarized the abundance of individuals (adults vs juveniles) across different activities. This test revealed a significant association between age group and activity type. To further explore this association, we calculated standardized residuals, which indicated which specific activity types were most strongly associated with each age group. We use the standard Adjusted Residuals threshold of 1.96 to -1.96 to determine whether the observed frequencies significantly deviated from expected values (MacDonald and Gardner 2000, García-Pérez and Núñez-Antón 2003), allowing us to identify the activities that were overrepresented or underrepresented for each age group. All analyses were performed in R version 4.2.1 (R Core Team 2023).

## Results

We obtained a total of 2,197 observation events, with 68.3% corresponding to flight activity, 14.7% to feeding, 13.1% to resting, and 3.2% to Comfort. For adult individuals, the highest activity peaks occurred between 7:00 - 7:29h and 15:00 - 15:29h. Juveniles exhibited three activity peaks: from 6:30 - 7:59h, 9:30 - 11:29h, and finally from 14:30 - 16:59h.

Human fishing activity was exclusively concentrated in the mornings between 7:00 – 9:00h. These activities usually require the fishermen to prepare their boats and equipment (hand lines, nets, traps, baits) in the early morning, sometimes before dawn. Then, they sail to the fishing grounds on boats that are made of wood and fiberglass or a mix of both. These bots are between 15 to 20 feet long and powered by outboard motors or paddles. Once in place, the fishermen check traps that were installed days before and set new traps with or without bait. Additionally, they fish using hand lines or nets. In El Cabo de la Vela the fishing activities were performed by 1 or 2 individuals from the local community. A maximum of three boats per fishing site were observed at the same time.

We performed several statistical tests to compare the start time and cumulative time spent on various activities between adults and juveniles, focusing on both circular and raw data.

### Watson’s Two-Sample Test of Homogeneity

We applied Watson’s two-sample test to compare the circular distributions of activity timing between adults and juveniles. The results indicated a significant difference in the circular patterns of activity between the two groups (Watson, p <0.001, Table 1), confirming that the distributions of activity times were not homogeneous. The analysis revealed significant differences in the timing and patterns of activities between adults and juveniles, particularly in resting and flight times. Adults and juveniles also differ in how they allocate their time across activities, especially in feeding and flight, as indicated by the Fisher’s exact test and residual analysis. The circular analysis confirmed that activity timing patterns differ significantly between the two age groups

**Table 1.** Circular Descriptive Statistics and Results of Watson’s Two-Sample Test of Homogeneity for Activity Patterns of Adult and Juvenile Brown Pelicans. The upper section of the table presents the circular descriptive statistics for the five activity categories: All Activities, Comfort, Resting, Feeding, and Flight. The mean activity time (in hours), standard deviation (SD), and circular concentration coefficient (Rho, ρ) are provided for both adults and juveniles. The Rho (ρ) statistic indicates the degree of clustering of activity times around the mean, with values closer to 1 representing higher concentration and values closer to 0 indicating higher dispersion. The lower section displays the results of Watson’s Two-Sample Test of Homogeneity, which was conducted to compare the circular distributions of activity times between adults and juveniles. Test statistics and p-values are reported for each activity. Significant differences (p < 0.001) were found for all activities, indicating that adults and juveniles exhibit significantly different activity timing patterns across all behavior categories.

| Circular descriptives |  |  |  |  |  |  |
| --- | --- | --- | --- | --- | --- | --- |
| Variable | Adults |  |  | Juveniles |  |  |
| | Mean | SD | Rho ($\rho$) | Mean | SD | Rho ($\rho$) |
| All activities | 11.050 | 1.039 | 0.582 | 11.670 | 0.912 | 0.659 |
| Comfort | 9.720 | 1.000 | 0.606 | 11.763 | 1.181 | 0.497 |
| Resting | 10.680 | 1.024 | 0.591 | 11.860 | 0.880 | 0.678 |
| Feeding | 11.909 | 1.038 | 0.583 | 11.443 | 0.870 | 0.684 |
| Flight | 10.951 | 1.033 | 0.586 | 11.453 | 0.902 | 0.665 |

**Table 1. Circular Descriptive Statistics and Results of Watson's Two-Sample Test of** **Homogeneity for Activity Patterns of Adult and Juvenile Brown Pelicans.**
| Watson's Two-Sample Test of Homogeneity |  |  |
| --- | --- | --- |
| Variable | Test statistic | P-value |
| All activities | 2.462 | < **0.001** |
| Comfort | 0.288 | < **0.01** |
| Resting | 0.464 | < **0.001** |
| Feeding | 0.388 | < **0.001** |
| Flight | 1.722 | < **0.001** |

### Start time

When analyzing the overall start times of all activities combined, both adults and juveniles showed non-normal distributions (Shapiro-Wilk, p < 0.001, Table 2). The Mann-Whitney U test confirmed a significant difference in activity timing between the two groups (Mann-Whitney U, p = < 0.001), indicating that juveniles and adults do not initiate activities at the same time (Fig 2).

**Table 2.**
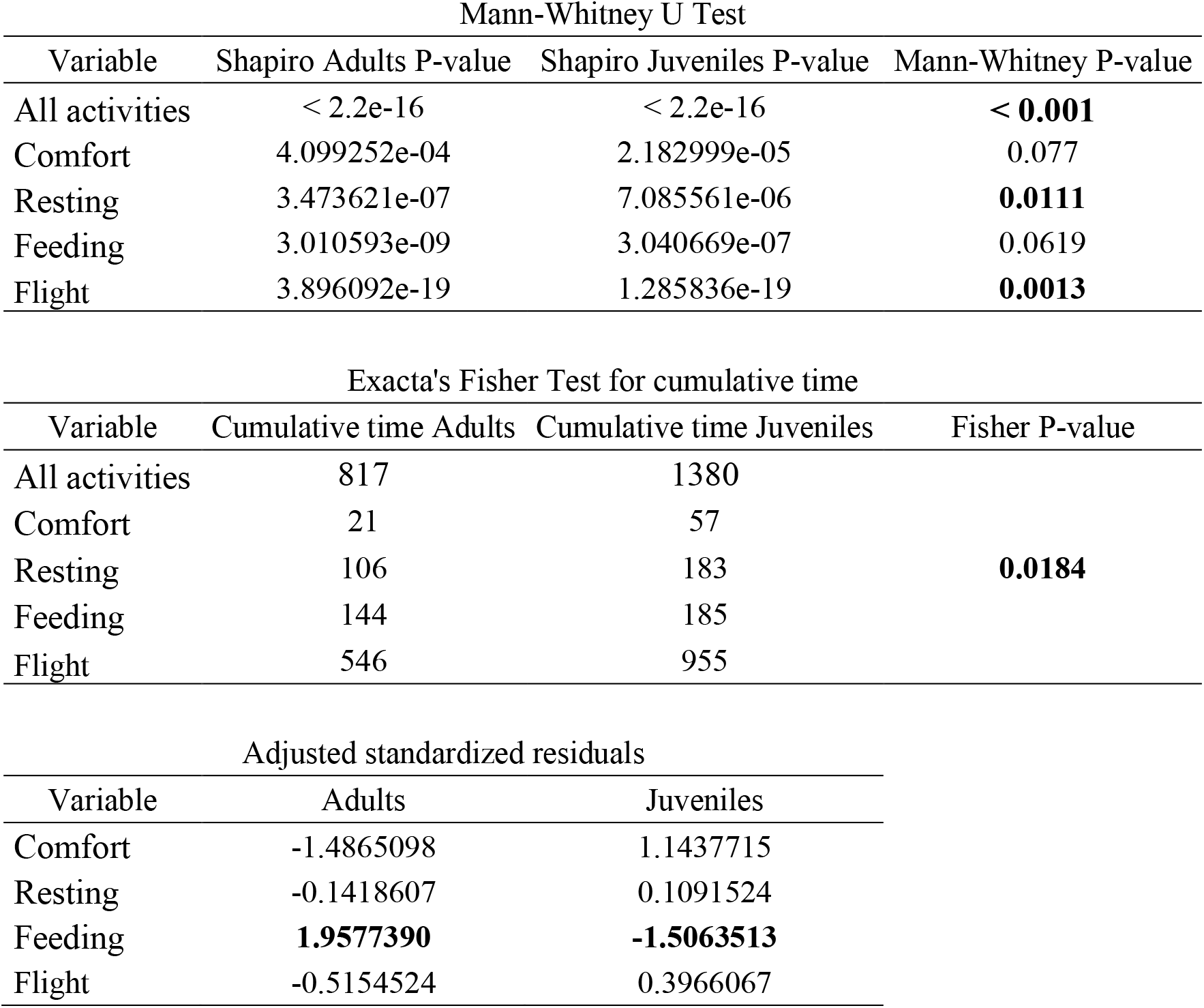
Mann-Whitney U Test for Start Time and Exact Fisher’s Test for Cumulative Time in Adult and Juvenile Brown Pelicans. The upper section of the table presents the results of the Mann-Whitney U test for comparing the start times of activities between adult and juvenile pelicans. The Shapiro-Wilk test was used to assess the normality of activity start times for both groups, with p-values indicating non-normal distributions (p < 0.001 for all variables). Significant differences in start times between adults and juveniles were found for Resting (p = 0.0111) and Flight (p = 0.0013), while Comfort and Feeding did not show statistically significant differences. The lower section shows the results of Exact Fisher’s test for cumulative time spent on activities by adults and juveniles. Juveniles spent significantly more cumulative time across all activities compared to adults (Fisher p = 0.0184), with marked differences particularly in Flight. Adjusted standardized residuals highlight overrepresented and underrepresented activity categories for each age group. Juveniles were overrepresented in Comfort activities (residual = 1.14) and underrepresented in Feeding (−1.51), while adults showed the opposite trend, being overrepresented in Feeding (1.96) and underrepresented in Comfort (−1.48). These results reflect significant differences in time allocation between the two age groups, indicating divergent activity strategies.

| Mann-Whitney U Test |  |  |  |
| --- | --- | --- | --- |
| Variable | Shapiro Adults P-value | Shapiro Juveniles P-value | Mann-Whitney P-value |
| All activities | < 2.2e-16 | < 2.2e-16 | **< 0.001** |
| Comfort | 4.099252e-04 | 2.182999e-05 | 0.077 |
| Resting | 3.473621e-07 | 7.085561e-06 | **0.0111** |
| Feeding | 3.010593e-09 | 3.040669e-07 | 0.0619 |
| Flight | 3.896092e-19 | 1.285836e-19 | **0.0013** |

| Exacta's Fisher Test for cumulative time |  |  |  |
| --- | --- | --- | --- |
| Variable | Cumulative time Adults | Cumulative time Juveniles | Fisher P-value |
| All activities | 817 | 1380 |  |
| Comfort | 21 | 57 |  |
| Resting | 106 | 183 | **0.0184** |
| Feeding | 144 | 185 |  |
| Flight | 546 | 955 |  |

**Table 2. Mann-Whitney U Test for Start Time and Exact Fisher's Test for Cumulative** **Time in Adult and Juvenile Brown Pelicans.**
| Adjusted standardized residuals |  |  |
| --- | --- | --- |
| Variable | Adults | Juveniles |
| Comfort | -1.4865098 | 1.1437715 |
| Resting | -0.1418607 | 0.1091524 |
| Feeding | **1.9577390** | **-1.5063513** |
| Flight | -0.5154524 | 0.3966067 |

**Figure 2.**
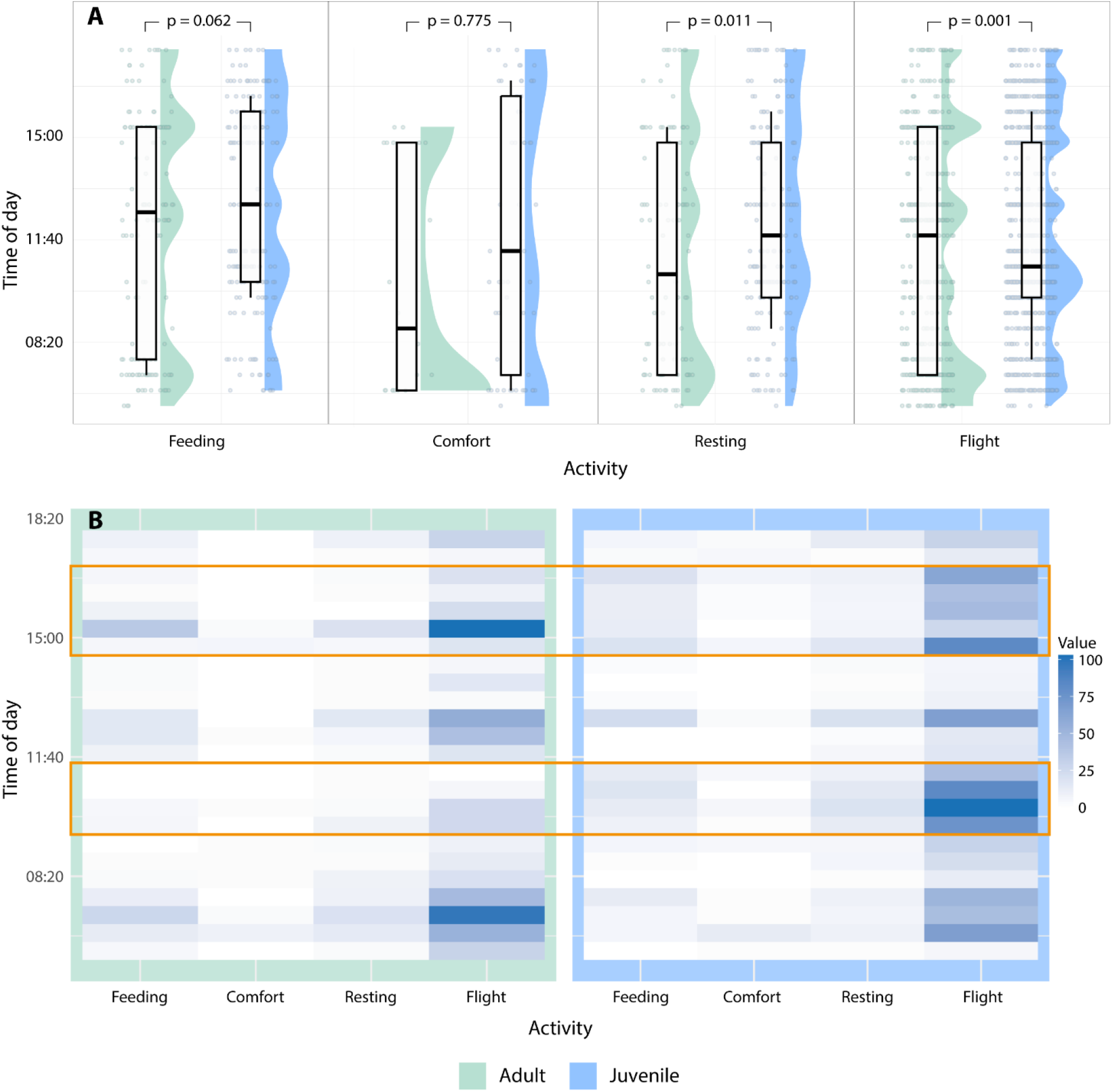
Comparison of Activity Patterns Between Juvenile and Adult Brown Pelicans. A) Raincloud plots showing the distribution of activity start times for four categories: Feeding, Comfort, Resting, and Flight, for both adults (green) and juveniles (blue). Each activity is represented with violin plots showing the distribution of data, boxplots indicating the interquartile ranges, and jittered individual data points. The statistical significance of differences between adults and juveniles for each activity is indicated by the p-values at the top of each plot. Significant differences are observed for Resting (p = 0.011) and Flight (p = 0.001), while no significant difference is observed for Feeding (p = 0.062) and Comfort (p = 0.775). B) Heatmap displaying the intensity of activity (value scale on the right) over the time of day for adults (left panel) and juveniles (right panel). Each row represents a time interval, and each column represents an activity type. Darker shades indicate higher activity levels. Orange areas represent a major contrast in the heatmap between adults and juveniles. Adults show prominent activity peaks for Resting and Flight around 15:00, while juveniles exhibit a similar flight peak but maintain higher flight activity earlier in the day (around 8:20).

Correspondingly, the Shapiro-Wilk test to assess normality in each activity group indicated that none of the distributions followed a normal pattern (p-values < 0.001 for all groups and activities). Consequently, we used the Mann-Whitney U to compare the timing of activities between adults and juveniles. For Resting, the test revealed a significant difference in the timing between the two groups (Mann-Whitney U, p = 0.011, Table 2), with juveniles starting their rest periods later than adults. For Flight, we also observed a significant difference (Mann-Whitney U, p = 0.001), suggesting that juveniles and adults engage in flight at different times (Fig 3). However, there were no significant differences in the timing of Feeding and Comfort activities (Mann-Whitney U, p = 0.061 and p = 0.77, respectively).

**Figure 3.**
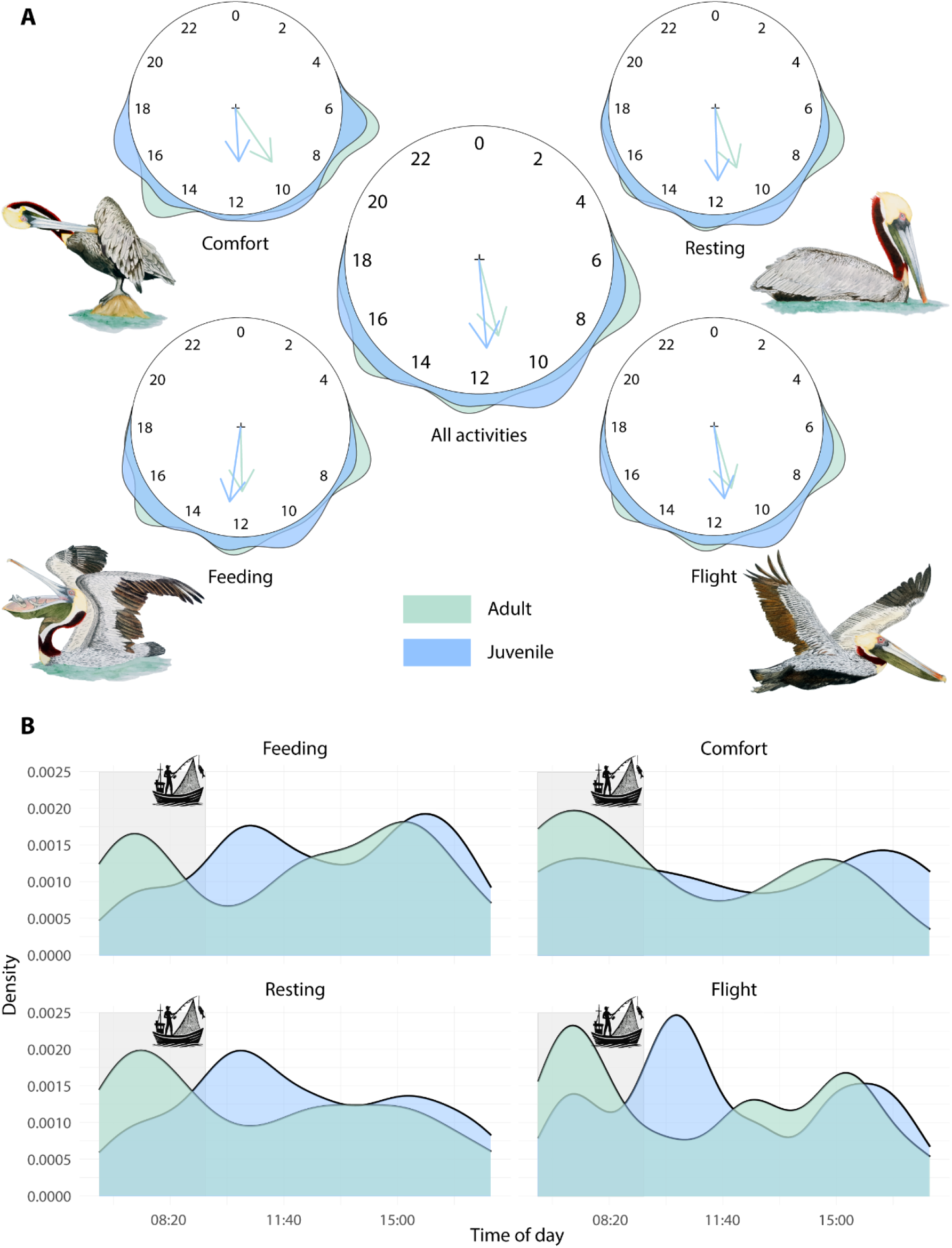
Circular and Density Plots of Brown Pelican Activity Patterns by Age Group. A) Circular plots representing the distribution of activity times for Comfort, Resting, Feeding, Flight, and All Activities combined, for adult (green) and juvenile (blue) Brown Pelicans. The direction of the arrows (ρ) indicates the mean activity time, while the length of the arrows reflects the concentration of the activity around that time. Juveniles tend to show more dispersed activity patterns across all categories compared to adults, with notable differences in the timing of Comfort and Flight activities. B) Density plots illustrating the distribution of activity occurrences over time for both adults (green) and juveniles (blue) across Feeding, Comfort, Resting, and Flight activities. The gray shade areas represent the presence of anthropogenic activity, particularly fishing boats, which influences the activity peaks. Adults generally show higher activity peaks earlier in the day which coincides with peaks in human activity. Feeding and Flight activities in juveniles are more prolonged throughout the day compared to adults, who concentrate their activities in shorter, more defined time windows. Pelican draws were made by Gabriela Serrato-Rivera ©.

### Cumulative Time Spent on Activities

We used Fisher’s exact test to examine the association between age group and activity type based on the cumulative time spent. The test revealed a significant difference between adults and juveniles (Fisher, p = 0.018), particularly in the Feeding category, where adults invested more time compared to juveniles. Conversely, juveniles spent considerably more time in Flight than adults. The Resting and Comfort activities did not show significant differences.

To further investigate which activity categories contributed most to the differences observed in the Fisher test, we performed a residual analysis. For Feeding, both adults and juveniles showed deviations from the expected values, with adults contributing more to the association (standardized residual = 1.96) and juveniles contributing less (−1.51) (Table 2). In contrast, no significant residuals were observed for the Resting and Flight activities, although the flight was more frequent among juveniles.

## Discussion

The activity patterns of juvenile and adult Brown Pelicans (*Pelecanus occidentalis*) studied at Cabo de la Vela, Colombia, revealed significant differences between age groups. These differences likely result from a combination of competitive dynamics, experience-based behavioral adaptation, and anthropogenic disturbances.

### Competition for Resources

Our findings suggest that adult seabirds may actively exclude juveniles from prime foraging areas, as evidenced by the temporal segregation of activity peaks. Adult pelicans exhibited their highest activity peaks in the early morning and late afternoon, while juveniles showed different peaks throughout the day. This temporal segregation may indicate competitive exclusion, a behavior seen in other seabird species (Fayet et al. 2015, Pettex et al. 2019). For instance, adult wandering albatrosses exhibit greater site fidelity and foraging efficiency than juveniles, leading to the displacement of juveniles to less productive areas (Grecian et al. 2018). Similarly, in Brown Pelicans, adults may occupy the best foraging locations during peak times, forcing juveniles to forage in suboptimal locations. This type of exclusion has also been documented in other seabirds like the Northern Gannet, where younger birds face competitive disadvantages in foraging success (Bodey et al. 2014).

One key behavior to consider in this context is the time spent flying, which is directly related to foraging in Brown Pelicans. As plunge divers, Brown Pelicans spend significant portions of their foraging time in flight, scanning the water’s surface for prey (Orians 1969, Walter et al. 2013). Therefore, the observed high flight activity in adults likely reflects their active searching and foraging during prime feeding times. In contrast, juveniles, which were observed to spend more cumulative time flying than adults, might be compensating for their lower foraging efficiency by increasing their flight time, possibly covering greater distances or needing more attempts to successfully catch prey (Orians 1969).

The association between flight and feeding supports the idea that the flight activity seen in both adults and juveniles is primarily linked to resource acquisition. In adults, this connection further strengthens the hypothesis of competitive exclusion: adults maximize foraging success during prime periods of the day, thus reducing juveniles’ opportunities for food acquisition. Juveniles, on the other hand, likely increase their flight time as a compensatory behavior due to their inferior foraging skills, a pattern that aligns with observations in other seabirds like the Northern Gannet (Bodey et al. 2014). The implications of this exclusion are critical, as access to sufficient food during early life stages can influence juvenile survival and long-term population dynamics.

### Behavioral Adaptation and Experience

In contrast to the hypothesis of competitive exclusion, our data also suggests that juveniles adjust their behavior to avoid direct competition with adults, suggesting behavioral adaptation. Juveniles engaged in flight and foraging activities at different times of the day, particularly in the mid-morning and early afternoon when adult activity was lower. This adaptation is consistent with findings from studies on juvenile wandering albatrosses, which show that juveniles develop distinct movement patterns and strategies to minimize competition with more experienced adults (de Grissac et al. 2016).

Furthermore, the “experience-based adaptation” hypothesis explains how juveniles may improve their foraging efficiency through increased flight activity over time. Juvenile Brown Pelicans, like other seabirds, must undergo a learning period where they refine their foraging techniques, including how to spot and capture prey from the air. Studies on wandering albatrosses show that juveniles gradually improve their foraging skills, becoming more efficient over time (Weimerskirch et al. 2006). In our study, juveniles exhibited longer flight times compared to adults, which may reflect their need to cover greater distances in search of food, or their inefficiency in identifying optimal foraging sites. This behavior, while less efficient, is an essential part of the learning process that will eventually lead to adult-level foraging efficiency.

### Anthropogenic Disturbance

Human activities, particularly fishing, have a well-documented impact on seabird foraging and movement behaviors. Our findings indicate a direct relationship in adult pelicans that altered their foraging behavior in response to the presence of fishing boats, which overlaps with the highest peak of activity. The availability of fishery discards provides an easily accessible food source, allowing seabirds to reduce their flight and search times (Bartumeus et al. 2010). The increased reliance on human-associated food sources can lead to shifts in activity patterns, potentially reducing competition between adults and juveniles as they adjust their behavior to exploit these new opportunities (Waugh et al. 2018, Afán et al. 2019).

Moreover, the presence of humans and fishing activities may also impose additional energetic costs on seabirds. When forced to relocate to avoid human disturbances, pelicans, particularly juveniles, may expend more energy, reducing their overall fitness and chances of survival. Studies on California Brown Pelicans have shown that human-induced disturbances, such as coastal development and fishing, can negatively impact foraging success and increase the energetic costs of flight and relocation (Lamb et al. 2018). The impact of such disturbances in our study site might exacerbate the already existing differences between adult and juvenile foraging behavior, further complicating the developmental trajectory of juvenile pelicans.

## Conclusions

The results of this study highlight the complex interplay between competition, experience, and human disturbance in shaping the activity patterns of Brown Pelicans. Juveniles and adults exhibit distinct behaviors, likely driven by competition for resources and differences in foraging efficiency. Over time, juveniles may adopt strategies to minimize direct competition with adults, developing their skills and foraging behaviors through experience. Additionally, human activities, such as fishing, play a significant role in altering natural behavior patterns, with implications for the conservation of pelican populations. This research underscores the importance of understanding intergenerational differences in activity patterns and the broader ecological and anthropogenic forces that influence these behaviors. Such insights are critical for the development of conservation strategies aimed at ensuring the survival and well-being of both juvenile and adult pelicans in the face of increasing environmental pressures.

## Acknowledgements

We would like to thank the Biology Program at Universidad El Bosque for their logistical support, which was essential for the successful completion of the fieldwork. We are deeply grateful to the community of the indigenous reserve Rancheria Kayuusipaa for their warm welcome and assistance during our stay. Thanks to Diego Medina Gaitán and Luis Serrano for their support in data collection, which significantly contributed to the success of this study, and thanks to Gabriela Serrato-Rivera for the beautiful drawings of pelican activities.

## Author contributions

DPC: Experiment design, conceptualization, data collection, writing and final review. JCRO: Conceptualization, data analysis, figures construction, writing original draft and final review. JDWV: Experiment design, data collection and final review.

## Funding

No financial support or budget was received for this work.

## Notes

https://dx.doi.org/10.6084/m9.figshare.27144603

### Competing Interest Statement

The authors have declared no competing interest.

